# The ALOG family members *OsG1L1* and *OsG1L2* regulate inflorescence branching in rice

**DOI:** 10.1101/2021.05.03.442460

**Authors:** Emanuela Franchini, Veronica M. Beretta, Israr Ud Din, Elia Lacchini, Lisa Van den Broeck, Rosangela Sozzani, Gregorio Orozco-Arroyo, Hélène Adam, Stefan Jouannic, Veronica Gregis, Martin M. Kater

## Abstract

The architecture of the rice inflorescence is an important determinant of seed yield. The length of the inflorescence and the number of branches are among the key factors determining the amount of spikelets, and thus seeds, that will develop. Especially the timing of the identity transition from indeterminate branch meristem to determinate spikelet meristem regulates the complexity of the inflorescence. In this context, the *ALOG* gene *TAWAWA1* (*TAW1*) has been shown to delay the transition to determinate spikelet development in rice. Recently, by combining precise laser microdissection of inflorescence meristems with RNA-seq we observed that two *ALOG* genes, *Oryza sativa OsG1-like 1* (*OsG1L1*) and *OsG1L2*, have an expression profile similar to *TAW1*. Here we report that *osg1l1* and *osg1l2* loss-of-function CRISPR mutants have similar phenotypes as the *taw1* mutant, suggesting that these genes might act on related pathways during inflorescence development. Transcriptome analysis of the *osg1l2* mutant suggested interactions of *OsG1L2* with other known inflorescence architecture regulators and the datasets were also used for the construction of a gene regulatory network (GRN) proposing interactions between genes potentially involved in controlling inflorescence development in rice.

The spatio-temporal expression profiling and phenotypical analysis of CRISPR loss-of-function mutants of the homeodomain-leucine zipper transcription factor gene *OsHOX14* suggest that the proposed GRN indeed serves as a valuable resource for the identification of new players involved in rice inflorescence development.

**One-sentence summary:** *OsG1L1* and *OsG1L2* control panicle architecture through delaying the transition from indeterminate branch- to determinate spikelet-meristem identity.

## INTRODUCTION

The inflorescences of land plants show a wealth of distinct architectures which evidences its importance for their reproductive success. The plant species or family-specific inflorescence shape depends on the activity and identity of meristems which determine the degree of branching and the number of flowers that will ultimately develop. Inflorescence meristems are defined as indeterminate since they continue to develop meristems in the axils of lateral organs, such as bracts. On the contrary, floral meristems are determinate as the development of the floral organs exhausts meristematic activity. In this sense, the formation of flowers can be seen as a developmental endpoint. An extreme example are tulips, where the apical meristem transforms into a floral meristem and forms one single apical flower. It is thus the identity transitions from indeterminate to determinate meristems that defines the complexity of an inflorescence (Hake, 2008).

*Oryza sativa*, commonly known as rice, develops a complex and determinate inflorescence, named panicle (Bommert et al., 2005; Han et al., 2014). Its architecture is established during early stages of rice reproductive development and it depends on the activity of different meristem types (Tanaka et al., 2013; Caselli et al., 2020). During the floral transition, the rice Shoot Apical Meristem (SAM) becomes Inflorescence Meristem (IM), also called rachis meristem. The IM gives rise to Primary Branch Meristems (PBMs) that produce Axillary meristems (AMs) which could differentiate into indeterminate Secondary Branch Meristems (SBMs) or determinate Spikelet Meristems (SMs). In the same way SBMs elongate and produce SMs. In rice, the SM develops three floral meristems (FMs), of which one will differentiate into one fertile floret, whereas the other two will develop into empty glumes (sterile lemmas); thus exhausting the pool of meristematic cells (Bommert et al., 2005; Han et al. 2014). Furthermore, the length of the rice inflorescence, and consequently the number of primary branches that can develop, is also determined by the timing of IM abortion.

Rice plants in which early transitions to spikelet meristem identity occur will have less complex panicles with fewer seeds in contrast to plants in which the transition is delayed. Among the genes that have been identified to control this transition are *ABERRANT PANICLE ORGANIZATION1 (APO1)* and *APO2* (Ikeda-Kawakatsu et al., 2012). Both are mainly expressed in IM and in BMs, where they also promote cellular proliferation. They are orthologs of the *Arabidopsis thaliana* genes *UNUSUAL FLORAL ORGANS* (*UFO*) and *LEAFY* (*LFY*), respectively. However, while the *UFO* and *LFY* promote floral identity, *APO1* and *APO2* repress the transition to determinate spikelet meristem formation (Ikeda-Kawakatsu, et al., 2009; Ikeda-Kawakatsu, et al., 2012). Recently, it was shown that LARGE2, a HECT-domain E3 ubiquitin ligase OsUPL2, interacts directly with APO1 and APO2 to modulate their stability. Genetic analysis of the *large2* mutant, which displays bigger panicles with more branches and seeds, confirmed that *LARGE2* functions in a common pathway with *APO1* and *APO2* (Huang et al., 2021).

*TAWAWA1* (*TAW1*)/*G1-LIKE 5* (*G1L5*) is another gene that promotes BM identity and suppresses SM specification by activating genes involved in the repression of floral transition (Yoshida et al., 2013). The dominant *taw1-D* gain-of-function mutant shows a delay in spikelet specification which results in increased branching and higher seed numbers. *TAW1* belongs to the *Arabidopsis LSH1 and Oryza G1* (*ALOG*) gene family, which includes fourteen *ALOG* genes in rice. The ALOG domain is highly conserved among land plants and evolutionary studies propose that it derived from the N-terminal DNA-binding domain of integrases that belong to the tyrosine recombinase superfamily which are encoded by a distinct type of DIRS1-like LTR retrotransposons that are found in several eukaryotes (Lyer & Aravind, 2012).

Recently, Harrop et al. (2016) used laser microdissection microscopy to specifically isolate different rice inflorescence meristem tissues for RNA-seq based expression profiling. Subsequent transcriptome analysis revealed that two *ALOG* genes, *OsG1L1* and *OsG1L2*, have similar expression profiles as *TAW1*. All three genes are highly expressed in the IM and subsequently, their expression gradually decreases in PBM, ePBM/AMs and SM. Phylogenetic analysis showed that *OsG1L1* and *OsG1L2* cluster both in subgroup A and are therefore more distantly related to *TAW1* which belongs to subgroup C (Li et al., 2019).

Here, we describe the functional analysis of *OsG1L1* and *OsG1L2* which evidenced that both genes play a similar role in rice inflorescence development. Like in the *taw1-3* and *TAW1* RNAi lines, both *osg1l1* and *osg1l2* CRISPR-Cas9 mutants showed a reduced branching phenotype and lower seed numbers. RNA-seq analysis of developing *g1l2* inflorescences was used to compute a gene regulatory network (GRN). Validation of the resulting network by a preliminary functional study of the homeodomain-leucine zipper transcription factor gene *OsHOX14* suggests that the proposed GRN could serve as a valid resource for the identification of genes controlling inflorescence development in rice.

## RESULTS

### *OsG1L1* and *OsG1L2* expression analysis

Previous transcriptome analysis of laser micro-dissected rice reproductive meristems, allowed identification of two *ALOG* genes, *OsG1L1* and *OsG1L2* that share a similar expression profile with the previously studied *TAW1* gene (Harrop et al., 2016; Yoshida et al. 2013; Figure 1A). These genes are highly expressed in the Inflorescence Meristem (IM), then mRNA abundance decreases in the Primary Branch Meristem (PBM), and a further reduction is observed in the elongated PBM with axillary meristem (ePBM/AM). In the Spikelet Meristem (SM) there is a slight increase in mRNA levels for all three genes.

**Figure 1.**
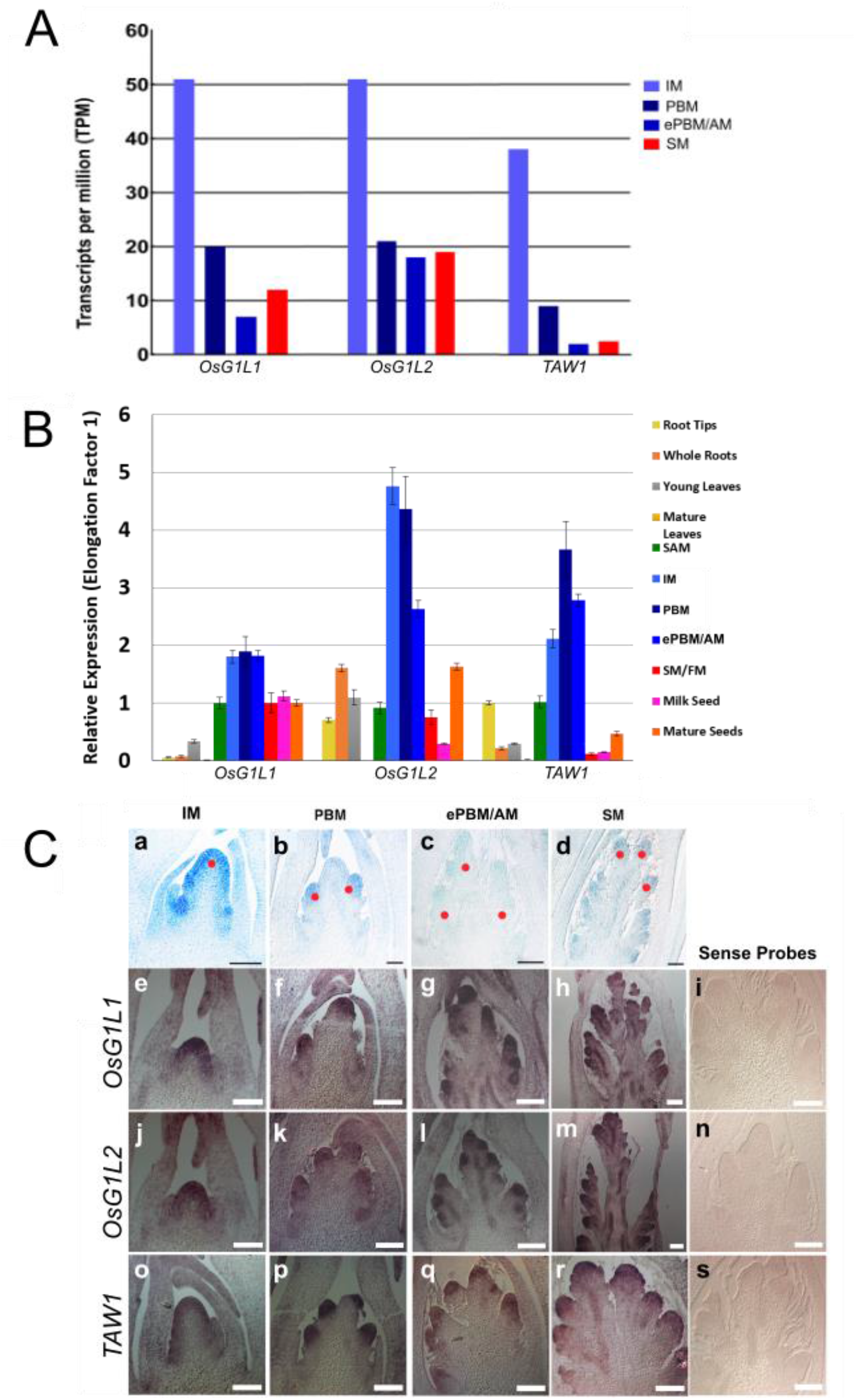
The *ALOG* family genes *OsG1L1, OsG1L2* and *TAW1* are highly expressed in inflorescence meristems. A, four reproductive meristem types sampled and analysed by RNA-seq (Harrop et al. 2016). Read counts are expressed in transcripts per million. B, qRT-PCR of *OsG1L1, OsG1L2* and *TAW1* across different tissues: vegetative tissues (roots and leaves), reproductive meristems enriched tissues manually dissected (IM, PBM, ePBM/AM, SM/FM) and seeds. C, Expression pattern of *OsG1L1, OsG1L2* and *TAW1* analysed by *in situ* hybridization at four developmental stages. Representation of four developmental stages that were analysed with reference to the meristem types indicated above the figure (a-d), where red dots indicate respectively the different meristematic tissues: IM (a), PBM (b), ePBM/AM (c) and SM (d). *OsG1L1* Antisense probe (e-h), *OsG1L1* sense negative control probe (i), *OsG1L2* antisense probe (j-m), *OsG1L2* sense negative control probe (n), *TAW1* antisense positive control probe (o-r), *TAW1* sense negative control probe (s). Scale bars represent 50 μm (a-c) and 100 μm (d-s). [Inflorescence Meristem (IM), Primary Branch Meristem (PBM), elongated PBM with Axillary Meristem (ePBM/AM), Spikelet Meristem (SM), Flower Meristem (FM)].

To determine in more detail the expression pattern of *OsG1L1* and *OsG1L2* and to confirm their expression in inflorescence meristems, we performed real-time PCR on different plant tissues such as, the root, the root tip (where the Root Apical Meristem (RAM) is localized), young and mature leaves, the Shoot Apical Meristem (SAM) and all reproductive meristems enriched tissues, like IM, PBM, ePBM/AM and SM/FM, and milk and mature seeds, respectively at 8 and 30 days after fertilization. Since the expression of *TAW1* was already described previously (Yoshida et al., 2013), it was used as a positive control. This analysis showed that all three genes are preferentially expressed in inflorescence tissues (IM, PBM, ePBM/AM and SM/FM) (Figure 1B).

RNA in situ hybridisation was performed to further investigate and compare the spatiotemporal expression of *OsG1L1, OsG1L2* and *TAW1* during different stages of panicle development. We designed for each of the three genes a specific digoxigenin-labelled RNA probe. This analysis revealed that *OsG1L1* and *OsG1L2* have a similar expression profile as *TAW1*. All three genes are expressed in the IM, PBM, ePBM/AM and SM/FM, suggesting *OsG1L1* and *OsG1L2* might have a similar functional role in the inflorescence meristems as *TAW1* (Figure 1C).

### Analysis of the *osg1l1* and *osg1l2* mutant phenotypes

To functionally characterize *OsG1L1* and *OsG1L2* CRISPR-Cas9 genome editing technology was used to generate mutations in these genes. A specific single-guide RNA (sgRNA) was designed for each gene and the two CRISPR-constructs (Miao et al. 2013) were used for *Agrobacterium tumefaciens* mediated transformation of rice embryonic calli. The two sgRNAs were designed to create indels in the ALOG domain.

The sgRNA designed for *OsG1L1* targeted the first exon at 397 bp from the ATG start site, whereas the sgRNA for *OsG1L2* was designed to target a region 131 bp downstream from the ATG start site (Supplemental Figure S1 A; Supplemental Figure S2 A). For *OsG1L1*, five T0 transgenic plants with different mutations at the sgRNA target site were obtained. Some of these mutations were in frame (3 and 6 bp deletions) and were not further analysed. Two independent transformants had a homozygous deletion of 2 bp (AG) at 145 bp from the translation start site. This mutation created a frameshift resulting in the formation of an aberrant protein, characterized by the disruption of the ALOG domain and of the putative nuclear localization signal (NLS). For these reasons, the obtained aberrant protein is most likely not functional (Supplemental Figure S1 B-C). We used for further analysis these two *osg1l1* mutants and in the T2 generation we obtained homozygous plants without the Cas9 encoding T-DNA insertion.

For the CRISPR-construct targeting *OsG1L2*, eighteen T0 transgenic rice plants were generated. All these plants had a similar frameshift mutation due to the insertion of a single base pair (A, C, G or T) at 148 bp from the start site. The A insertion leads to the formation of a premature stop codon (TGA), resulting in the production of a truncated protein of 49 amino acids (Supplemental Figure S2 B-C), lacking the ALOG domain and the putative NLS. The insertion of one of the other bases (C, G, T) leads to the formation of an out of frame reading frame resulting in a protein of 176 amino acids without the ALOG domain and the putative NLS (Supplemental Figure S2 C). We selected in the T1 generation two independent *osg1l2* mutant lines having an A or C insertion. In the T2 generation we obtained lines homozygous for these insertions and without the Cas9 encoding T-DNA insertion. For detailed phenotyping we used the line with the A insertion.

During the vegetative growth, the *osg1l1* and *osg1l2* mutants didn’t show any obvious phenotype (data not shown). Considering that the expression of *OsG1L1* and *OsG1L2* was predominantly restricted to inflorescence meristem tissues, a detailed phenotypic analysis of the panicle was performed using Panicle TRAit Phenotyping software (P-TRAP) (Figure 2; AL-Tam et al., 2013). To obtain a robust statistical analysis, at least 15 plants for each genotype were analysed. In particular we analysed 15 wild-type plants, 19 *osg1l1* mutants and 20 *osg1l2* mutants. After panicle imaging, the P-TRAP software quantified the traits related to the panicle architecture and seed numbers (Supplemental Table S1).

**Figure 2.**
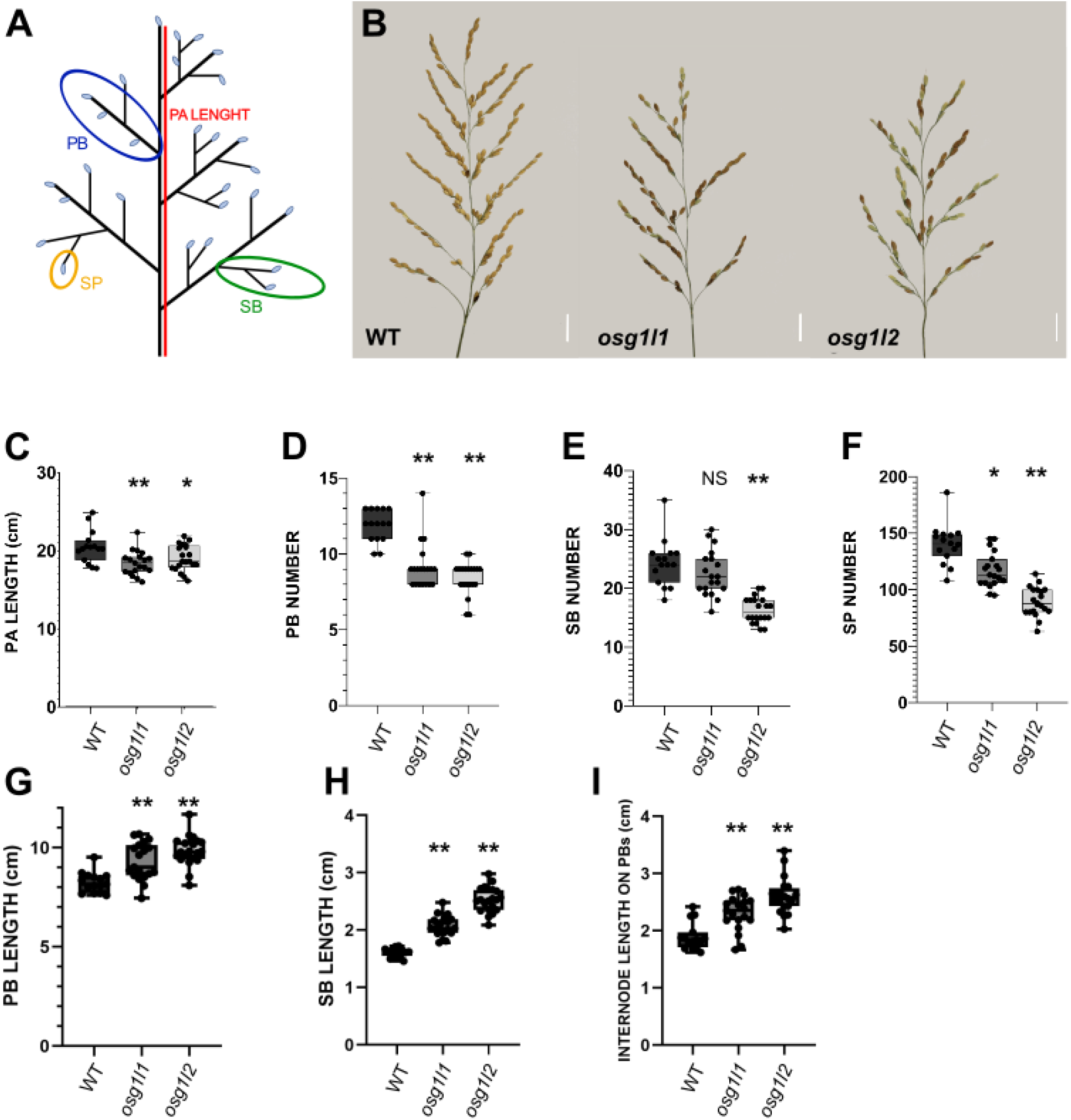
Phenotypical analysis of panicle architecture in wild type, *osg1l1* and *osg1l2* mutants. A, Schematic representation of panicle structure [PA= panicle; PB= primary branches; SB= secondary branches; SP= spikelets/seeds]. B, Main panicles of WT, *osg1l1* and *osg1l2* (2 cm scale bars). Graphs representing the comparison of: (C) Panicle (PA) Length, (D) Primary Branch (PB) number, (E) Secondary Branch (SB) number, (F) Spikelet/seed (SP) number in WT, *osg1l1* and *osg1l2* backgrounds. Graphical representation of the comparison between the: (G) length of Primary Branches (PB), (H) length of Secondary branches (SB) and (I) length of the internodes in PBs. One-Way ANOVA with Tukey test; **p<0,01; * p<0,05.

This analysis showed that *osg1l1* and *osg1l2* produced significantly shorter panicles than wild-type plants. Furthermore, their panicles developed fewer PBs, SBs and spikelets/seeds. In particular, the *osg1l1* mutants produced panicles that were on average 2 cm shorter than wild type and developed on average 3 PB and 30 spikelets less than wild-type plants. Interestingly, the number of SBs were not significantly different from wild type (Figure 2 C-F).

The *osg1l2* mutant plants produced panicles that were on average 1.5 cm shorter than wild type and developed on average 4 PBs, 8 SBs and 51 spikelets/seeds less than wild-type plants (Figure 2 C-F). This analysis confirmed a previous independent experiment in which PBs, SBs and spikelet/seed numbers were compared between wild-type and *osg1l2* plants with a different mutation (a C insertion at 148 bp from the ATG) (Supplemental Figure S3).

The P-TRAP analysis revealed also that both *osg1l1* and *osg1l2* mutant lines had longer PBs and SBs as well as longer internodes in PBs (Figure 2 G-I). In detail, the *osg1l1* mutant plants displayed PBs and SBs that were on average respectively 1 cm and 0.5 cm longer than those of wild-type rice plants. The internodes of PBs were on average 0.4 cm longer than in wild type. The *osg1l2* mutant plants instead produced PBs and SBs that were on average 1.6 cm and 1 cm longer than wild type, respectively. The internodes of PBs were on average 0.7 cm longer than wild type. Overall, both mutants showed similar aberrations in panicle architecture although the phenotype of the *osg1l2* mutant was more severe.

To investigate whether loss-of-function of *OsG1L1* and *OsG1L2* not only affected seed numbers but also seed size, the length, width and surface area of *osg1l1* and *osg1l2* mutant seeds were measured. For each genotype (*osg1l1, osg1l2* and wild type) a minimum of 100 seeds were analysed using the Smart Grain software (Tanabata et al., 2012; Supplemental Table S2). This analysis revealed that *osg1l1* and *osg1l2* mutants produced significantly smaller seeds than wild type. Both *osg1l1* and *osg1l2* seeds had smaller seed areas, where *osg1l1* seeds had a reduced length and width and *osg1l2* seeds only showed a reduction in seed width compared to wild type (Figure 3).

**Figure 3.**
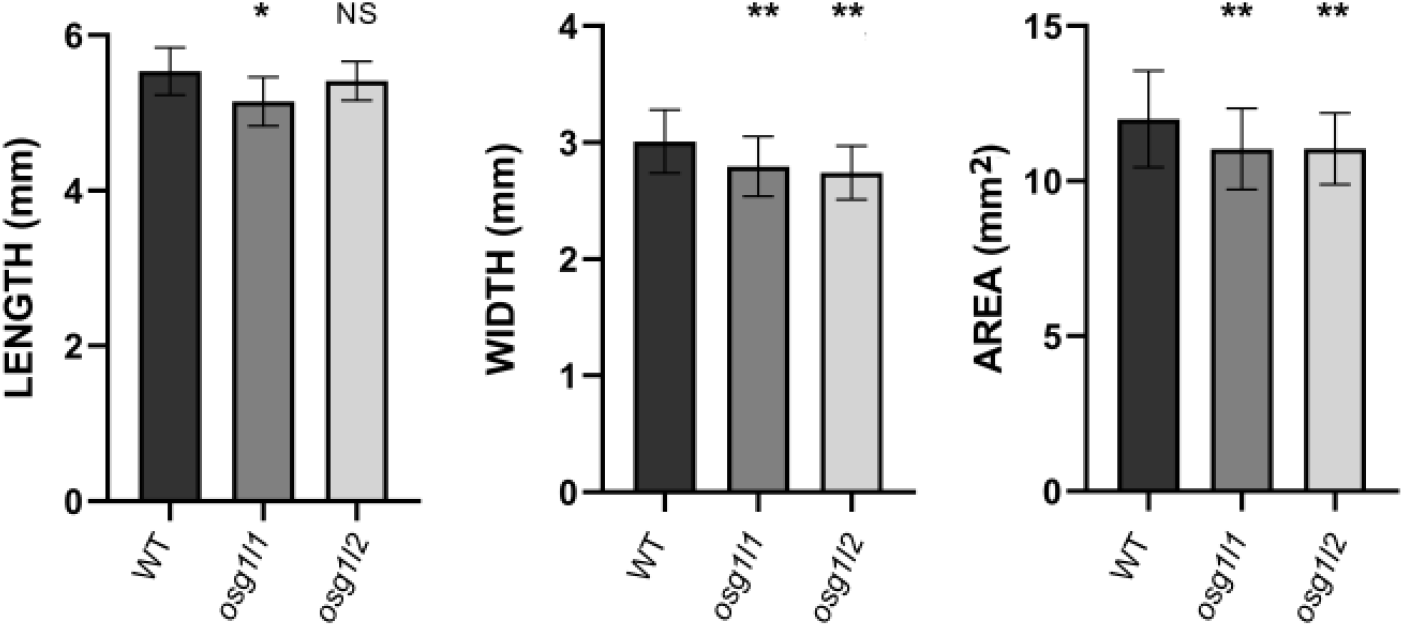
Size measurements of wild type, *osg1l1* and *osg1l2* seeds. Graphs resulting from the analysis of length, width and area of wild type (WT), *osg1l1* and *osg1l2* seeds using the Smart Grain software (Tanabata et al., 2012). 100 seeds for each genotype were analysed. Statistical One-Way ANOVA with Tukey test: **p<0,01; * p<0,05. NS = not significant.

### Transcriptome analysis of the *osg1l2* mutant at early stages of inflorescence development

The phenotypic analysis of the *osg1l1* and *osg1l2* mutants suggests that both genes may play a similar role during rice inflorescence development. Since inflorescence branching was more severely affected in the *osg1l2* mutant, this line was selected for RNA-seq transcriptome analysis to obtain deeper insights into the role this *ALOG* gene plays at the early stages of inflorescence development. Developing inflorescences of the wild type and the *osg1l2* mutant at early developmental stages enriched in PBMs and ePBM/AMs were manually dissected. Material for four biological replicates was obtained, each replicate consisting of 8 to 10 dissected inflorescences. Subsequently, RNA was extracted and used for Illumina sequencing.

The raw RNA-seq files were processed using the TuxNet interface (Spurney et al., 2019). Reads were cleaned, mapped on the *O. sativa* reference genome (IRGSP-1.0), normalized, and FPKMs (Fragment per kilobase of transcript per million mapped reads) were calculated (Supplemental Table S3). Finally, performing a pairwise differential expression analysis between the wild type and mutant, the TuxNet interface generated Differentially Expressed Gene (DEG) datasets (Spurney et al., 2019) (Supplemental Table S4).

After data processing, an average of approximately 13.000.000 reads for each replicate were obtained. The alignment of the reads to the reference genome resulted overall in > 97% coverage. Setting the log_2_ (Fold Change (FC)) equal to 1 and the q-value equal to 0.05, a total of 279 differentially expressed genes were identified, of which 152 were downregulated and 127 upregulated in the *osg1l2* mutant compared to the wild type. <u>Among</u> the upregulated genes in the *osg1l2* mutant, we identified *OsMADS37*, a MADS-box transcription factor encoding gene homologous to Arabidopsis *FLOWERING LOCUS C* (*FLC*) (Ruelens et al., 2013), *OsERF112*, an ERF/AP2 transcription factor (Nakano et al., 2006), and *OsG1L4*, another member of the *ALOG* gene family. Interestingly, also *EPIGENETIC SHORT PANICLE* (*OsESP*), a putative long-noncoding RNA whose overexpression leads to shorter and denser panicles, was upregulated in the *osg1l2* mutant (Luan et al., 2019). Moreover, the expression of genes involved in hormonal pathways was upregulated, such as *OsRR3*, an A-type response regulator that acts as a negative regulator of cytokinin signalling (Cheng et al., 2010). Among the upregulated genes there are several which encode for Zinc-finger transporter proteins and genes encoding for proteins containing a NB-ARC domain, which is associated to plant disease resistance (Van Ooijen et al., 2008) (Supplemental Table S4 and Table 1).

**Table 1.** Genes differentially expressed in the *osg1l2* mutant and involved in inflorescence development. For each gene is indicated the Gene Name (column one); the Gene ID (column two); the log2 Fold Change (log2 FC, column three); the Q-value (column four; a significance cut-off of0.05 is applied) and information related to their function (column 5).

| Gene name | Gene ID | log2 FC | Q-value | Putative function |
| --- | --- | --- | --- | --- |
| ***OsESP*** | Os01g0356951 | 6,55E+04 | 0,00245724 | Involved in the regulation of panicle architecture |
| ***OsMADS37*** | Os08g0531900 | 1,87E+00 | 0,00245724 | Homolog of *FLC* in rice |
| ***OsG1L4*** | Os04g0516200 | 1,34E+00 | 0,00245724 | Protein of unknown function DUF640 domain containing protein |
| ***OsERF112*** | Os12g0603300 | 1,09E+00 | 0,00245724 | Similar to AP2 domain containing protein |
| ***OSRR3*** | Os02g0830200 | 1,95E+00 | 0,00245724 | A-type response regulator involved in Cytokinin signaling |
| ***ONAC120*** | Os10g0477600 | -2,47E+00 | 0,00245724 | Similar to NAM / CUC2-like protein |
| ***OsTCP25*** | Os09g0521300 | -1,07E+00 | 0,0485909 | Similar to *TCP* family transcription factor |
| ***OsTB1*** | Os03g0706500 | -1,12E+00 | 0,00245724 | *TCP* family transcription factor, negative regulator of lateral branching |
| ***OsPP2C30*** | Os03g0268600 | -1,01E+00 | 0,00245724 | Similar to Protein phosphatase type 2C |
| ***OsMADS34/PAP2*** | Os03g0753100 | -1,13E+00 | 0,00245724 | MADS-box transcription factor, involved in inflorescence and spikelet development |
| ***OsRCN1*** | Os11g0152500 | -1,04E+00 | 0,00245724 | Putative phosphatidylethanolamine-binding protein, Rice *TFL1/CEN* homolog. Involved in the control of inflorescence architecture and in the repression of flowering |
| ***OsRCN4*** | Os04g0411400 | -1,48E+00 | 0,00839011 | Terminal flower 1-like protein |
| ***OsHOX14*** | Os07g0581700 | -1,53E+00 | 0,00456853 | Homeodomain-leucine zipper (HD-Zip) transcription factor, may be involved in the regulation of panicle development |
| ***OsCEP6*** | Os08g0475500 | -1,02E+00 | 0,00245724 | C-terminally encoded peptide, involved in the regulation of panicle and grain development |
| ***OsPP2C1*** | Os09g0325700 | -2,08E+00 | 0,00245724 | Protein phosphatase 2C. Involved in abiotic stress response and early panicle development |
| ***OsGATA7*** | Os10g0557600 | -1,49E+00 | 0,00245724 | GATA transcription factor. Involved in brassinosteroids-mediated growth regulation, panicle development and grain shape/number/weight/yield |
| ***OsIAA14*** | Os03g0797800 | -1,44E+00 | 0,00245724 | Protein belonging to the AUX/IAA protein family |

Among the downregulated genes, we found TFs like *ONAC120* that is similar to NAM/CUC-2 like protein (Ooka et al., 2003) and two genes belonging to the TCP family, *OsTCP25* and *OsTB1/FC1* (that henceforth will be referred to as *OsFC1*). Interestingly, *OsFC1* is already known to be a negative regulator of tillering and inflorescence development (Takeda et al., 2003; Cui et al., 2020). Notably, the transcription factors *OsMADS34/PAP2* and *OsGATA7*, which are known to be involved in inflorescence architecture establishment; and *OsHOX14*, which has been proposed to be involved in panicle development, were also downregulated (Gao et al. 2010; Kobayashi et al., 2012; Shao et al., 2018; Zhang et al., 2018). Additional downregulated genes that have been associated with inflorescence development are *OsPP2C1* (Protein phosphatase 2C), *OsRCN1* and *OsRCN4* (Putative phosphatidylethanolamine-binding protein and Rice TFL1/CEN homolog), and *OsCEP6* (C-terminally encoded peptide) (Nakagawa et al., 2002; Li et al., 2013; Sui et al., 2016). Moreover, we could also find genes involved in hormonal pathways like *OsIAA14*, belonging to the Aux/IAA family and involved in auxin-response (Jain et al., 2006) (Table 1 and Supplemental Table S4).

### Gene regulatory network inference predicts a functional role for *OsHOX14*

To identify major regulatory transcription factors underlying the inflorescence phenotype of *osg1l2*, we built a gene regulatory network (GRN) using a regression tree with random forest approach (Spurney et al., 2019). Specifically, we inferred causal relations between 15 identified differentially expressed transcription factors (TFs) (log_2_(FC) > 1 or < −1, q-value < 0.05) and 232 downstream genes in *osg1l2* mutant (log_2_(FC) > 1 or < −1, q-value < 0.05). The inferred network contained 79 genes, of which five TFs have more than 10 outgoing regulations, including *OsWRKY80* (Wu et al., 2004), *OsG1L2, OsMADS37* (Ruelens et al., 2013), *OsFC1* (Takeda et al., 2003; Cui et al., 2020), and *OsGATA7* (Zhang et al., 2018) (Figure 4). One of these major regulators is *OsG1L2*, regulating 16 downstream genes, several of which have been shown to be involved in rice inflorescence development (Figure 4 C). For example, *OsESP* is a putative long-noncoding RNA whose gain-of-function mutant leads to a short and denser panicle (Luan et al., 2019) and *OsJAZ4*, also known as *OsTIFY11b*, is a positive regulator of grain-size acting downstream of *TRIANGULAR HULL1* (*OsTH1*), another ALOG factor (Hakata et al., 2012; Wang et al., 2019). Interestingly, thirteen of the predicted OsG1L2 targets are upregulated in the *osg1l2* mutant, ten of which are predicted to be co-regulated by *OsFC1*, a gene known to negatively regulate branching (Takeda et al., 2003; Cui et al., 2020).

**Figure 4.**
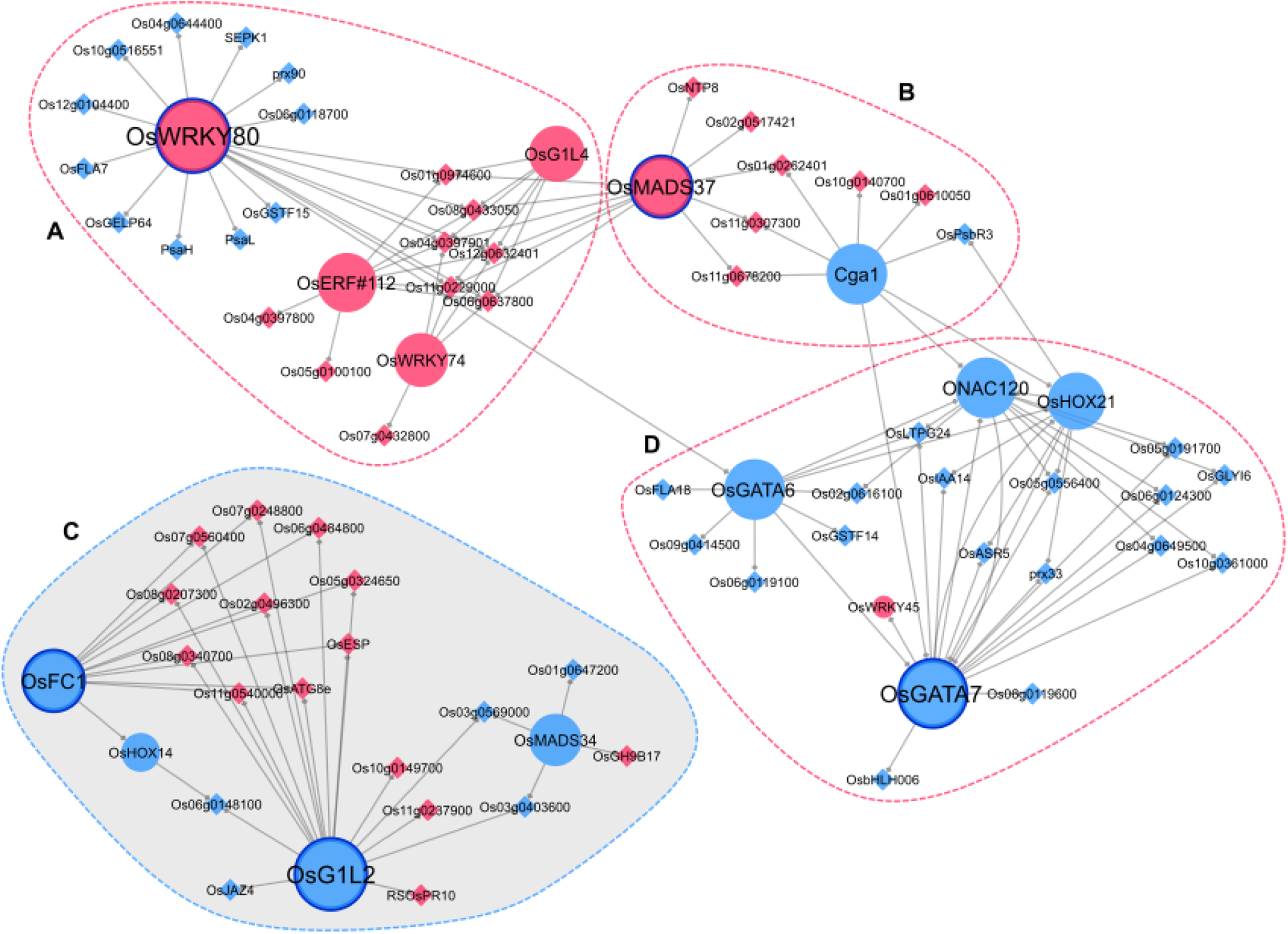
Graphic representation of the predicted Gene regulatory Network (GRN). Regulatory interactions were inferred using a regression tree with random forest approach. Transcription factors and other genes are represented in circles and diamonds, respectively. The interactions are represented by a diamond arrow. Upregulated and downregulated genes are highlighted in magenta and light blue respectively. Circled with a dotted line, there are the four major subclusters (A to D) in which the network can be divided. Circled in blue with a continuous line there are the five major TFs of the GRN: *OsWRKY80, OsG1L2, OsMADS37, OsFC1*, and *OsGATA7*. The module containing *OsG1L2* is highlighted in grey (C).

To identify regulatory subclusters within the *osg1l2* network, we clustered the network genes into different modules with the Cytoscape plugin clusterMaker2 (see materials and methods) (Figure 4). A total of four modules were identified, one smaller module of 10 genes (Figure 4 B), and three larger modules of 24, 23 and 22 genes (Figure 4 A, C, D). The smaller module contains *CYTOKININ-RESPONSIVE GATA TRANSCRIPTION FACTOR1* (*OsCGA1*), of which constitutive overexpression reduced grain filling (Hudson et al., 2013) and *OsMADS37*, the closest homologues of Arabidopsis *FLOWERING LOCUS C* (Shrestha et al., 2014) (Figure 4 B). One of the largest modules contains two *GATA* TFs, including *OsGATA7*, which influences architecture and grain shape and of which CRISPR/Cas9 lines show a similar phenotype as *osg1l2* (Zhang et al., 2018) (Figure 4 D). As we were interested in the regulatory interactions underlying the *osg1l2* mutant phenotype, we focused on module C, which contains four TFs: *OsG1L2, OsFC1, OsMADS34*, and *OsHOX14*. Interestingly, *OsMADS34* has been shown to be necessary for correct inflorescence development, further emphasizing the functional importance of this regulatory module (Kobayashi et al., 2012). *OsHOX14* can form heterodimers with *OsHOX12*, a gene that regulates panicle exertion (Gao et al., 2016). Overall, our network analysis allowed for the identification of many interesting candidates, of which several have been described in the context of panicle development, and indicated putative targets of OsG1L2 and other major TFs, giving also hints of which genes might be involved in the same developmental pathway regulating a similar set of genes.

### Validation of GRNs by qRT-PCR

RT-qPCR experiments were carried out to validate some of the deregulated genes of subcluster C of the predicted GRN (Figure 4). In particular, we focused on those genes that were already proposed to be involved in inflorescence development, such as *OsHOX14, OsMADS34, OsFC1* and *OsESP* (Takeda et al., 2003; Gao et al. 2010; Shao et al., 2018; Luan et al., 2019). *Os03g0569000* was also analysed because, according to the GRN, it was a common target of *OsMADS34* and *OsG1L2*. The qRT-PCR was performed using three biological replicates of developing inflorescences enriched in PBMs and ePBM/AMs of wild-type and the *osg1l2* mutant. Furthermore, the expression of the selected genes was also analysed in the *osg1l1* mutant, to determine if these genes might be regulated by both *ALOG* genes.

As shown in Figure 5 (A-E), the qRT-PCR confirmed the downregulation of *OsHOX14, OsMADS34, OsFC1* and *Os03g0569000* and the upregulation of *OsESP* in *osg1l2* mutant compared to WT. The expression analysis performed on the selected genes in the *osg1l1* mutant highlighted that *OsHOX14, OsMADS34, OsFC1* and *Os03g0569000* were also downregulated in the *osg1l1* mutant (Figure 5B, 5C and 5D); whereas the expression level of *OsESP* was not significantly different from wild-type inflorescences (Figure 5E). Overall, these results suggest that the analysed genes within the OsG1L2-containing subcluster C (Figure 4) might have a genetic interaction as predicted by the GRN.

**Figure 5.**
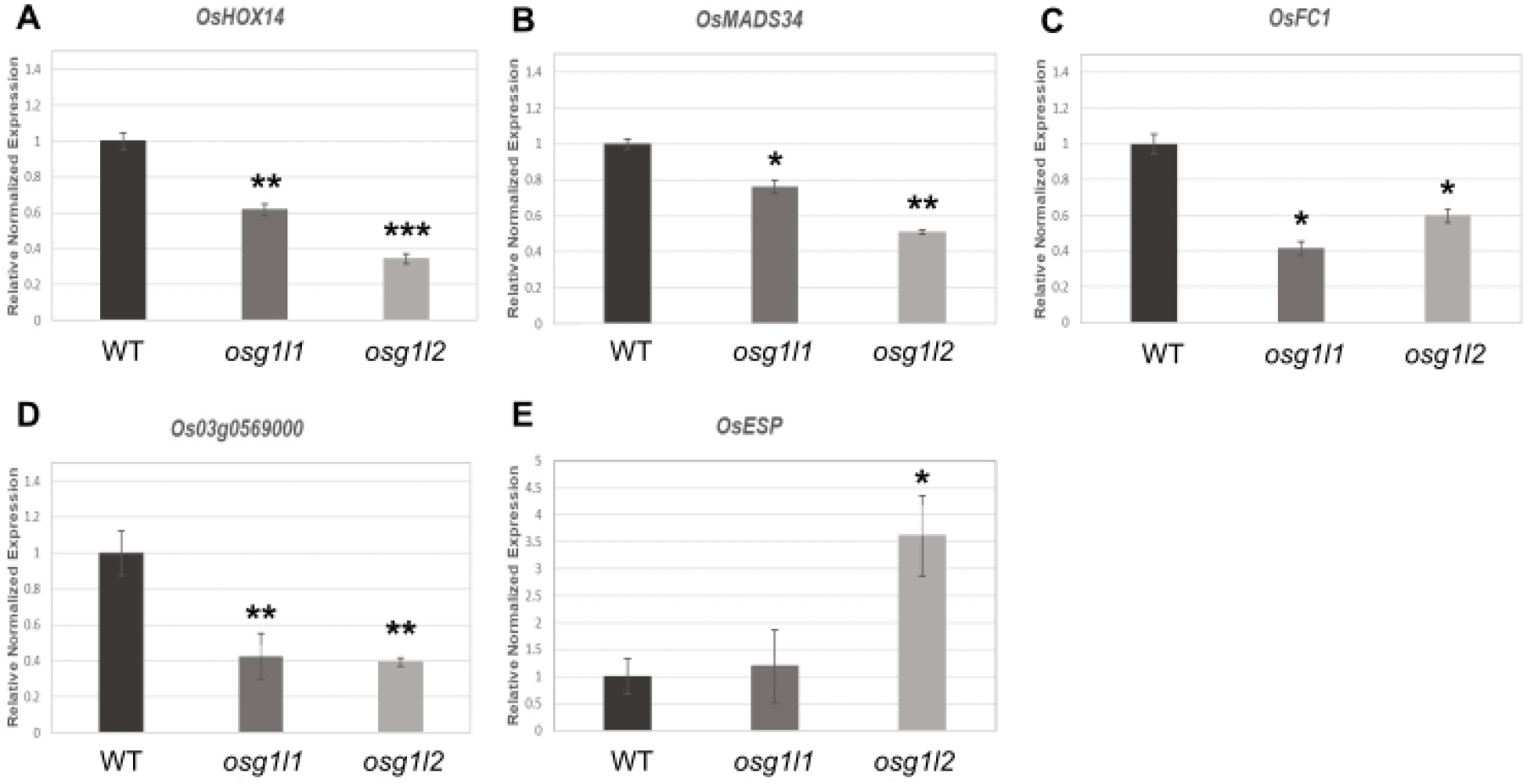
Expression analysis of subcluster C genes in the *osg1l1* and *osg1l2* mutants. Expression analysis of *OsHOX14* (A), *OsMADS34* (B), *OsFC1* (C), *Os03g0569000* (D), *OsESP* (E) by quantitative real-time PCR in wild type (WT), *osg1l1* and *osg1l2* mutants. Expression of *OsHOX14, OsMADS34, OsFC1* and *Os03g0569000* was normalized to that of *Elongation Factor 1* and the expression level of wild type was set to 1. The asterisks indicate: *** p< 0,001; **p<0,01; * p<0,05, student’s t-test.

### *Oshox14* mutant exhibiting a branching phenotype

The GRN subcluster C, that contains *OsG1L2*, shows as major components *OsMADS34, OsFC1* and *OsHOX14. OsMADS34* and *OsFC1* have been intensively studied for their role in inflorescence development (Takeda et al., 2003; Gao et al. 2010) but for *OsHOX14* only an overexpression study has been reported (Shao et al., 2018). Therefore, we selected *OsHOX14* for further functional studies to validate the GRN predicted involvement of this gene in inflorescence development.

*OsHOX14* is a member of the homeodomain-leucine zipper (HD-Zip) transcription factor family and it is the rice orthologue of barley *HvHox2* that, together with its recently diverged paralogue *HvVrs1*, are responsible for either the suppression or establishment of the barley lateral sterile spikelets (Sakuma et al., 2010). Indeed, *OsHOX14*, like *OsG1L1* and *OsG1L2* was shown to be expressed in the inflorescence meristem tissues (PBM, ePBM/AM, SM) (Harrop et al., 2016). The spatiotemporal expression of *OsHOX14* throughout early panicle development was assessed with an *in situ* hybridization experiment. This TF resulted to be expressed in PBM, ePBM/AM and SM/FM, suggesting a putative role of this gene during reproductive meristems establishment (Figure 6 A-C). We generated a knock-out mutant line for *OsHOX14* using the CRISPR-Cas9 genome editing system (Miao et al., 2013). A specific sgRNA was designed to target the first exon of the *OsHOX14* gene (Supplemental Figure S4A). T0 transgenic plants were selected and genotyped. In the T1 generation, three different mutant lines were obtained, having respectively, a homozygous G deletion, a homozygous C insertion, and a homozygous CG deletion and T insertion at 7 bp, 11 bp, and 6 bp downstream the start site. In all three cases, the different mutations led to a frameshift in the coding sequence and the formation of a premature stop codon, which resulted in the formation of a protein consisting of 22, 27, and 22 amino acids, respectively (Supplemental Figure S4 B-C). To evaluate the inflorescence phenotype of *oshox14*, a comparative analysis was performed on panicles belonging to 5 wild-type and 5 T1 *oshox14* mutant plants (1 plant carrying the homozygous C insertion mutation, 1 plant carrying the CG deletion and T insertion mutation and 3 plants carrying the G deletion mutation). This analysis revealed that the *oshox14* mutants developed panicles with less PBs and spikelets/seeds than wild-type plants. The number of SBs was not significantly different from wild-type. In particular, the *oshox14* mutant plants produced panicles that developed on average 3 PBs and 20 spikelets less than the wild-type (Figure 6 D-G). Overall, this analysis shows that *OsHOX14* plays a role in inflorescence branching as predicted by the GRN.

**Figure 6.**
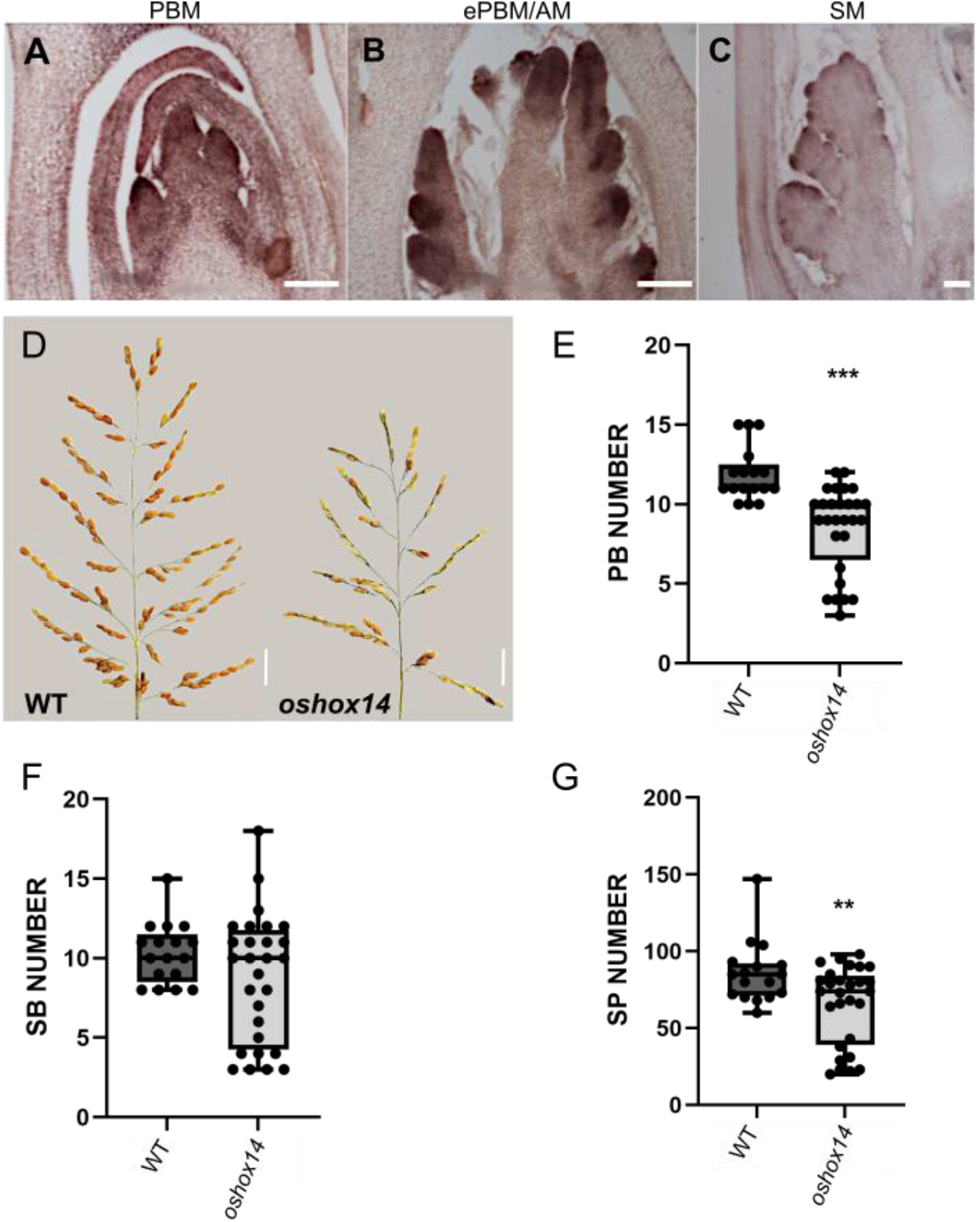
Expression and function analysis of *OsHOX14*. *In situ* analysis of three developmental stages: A, PBM; B, ePBM/AM and C, SM. Scale bars represent 50 μm (C) and 100 μm (A-B). [Primary Branch Meristem (PBM), elongated PBM with Axillary Meristem (ePBM/AM), Spikelet Meristem (SM)]. D-G, Phenotypical analysis of panicle architecture in wild type and *oshox14* mutant. D, main panicles of wild type (WT) and *oshox14* (2 cm scale bars). Graphs representing the comparison of: E, Primary Branch (PB) number; F, Secondary Branch (SB) number; G, Spikelet/seed (SP) number in WT and *oshox14* backgrounds. *** = p< 0,001; ** = p<0,01; * = p<0,05, Student’s t-test.

## DISCUSSION

Inflorescence architecture is a key agronomic trait that influences plant yield and its development is finely regulated by genes involved in meristem identity specification and in the control of the transition from indeterminate to determinate growth. Therefore, identifying genes involved in inflorescence development promises to contribute to improved crop yield through breeding and biotechnological approaches.

In this study, we functionally characterized *OsG1L1* and *OsG1L2*, two rice genes belonging to the ALOG gene family. These two genes are likely to play an important role in the regulation of inflorescence architecture specification acting as positive regulators of primary and secondary branch development. *OsG1L1* and *OsG1L2* were shown to be preferentially expressed in the reproductive meristems, in a pattern similar to *TAW1*, a gene already known to be involved in the development of the rice inflorescence by promoting indeterminate meristem identity (Yoshida et al 2013). Phenotypical analysis of the *osg1l1* and *osg1l2* mutant inflorescences showed that the two single mutants developed shorter panicles with fewer spikelets when compared to wild type. Furthermore, the mutant seeds were also shown to be smaller.

The observed aberrations in inflorescence architecture of the *osg1l2* mutant and to less extent of the *osg1l1* mutants indicate that they might be involved, like *TAW1*, in the regulation of the identity transition from BM to SM.

However, the DEG list obtained from our RNA-seq analysis of the *osg1l2* mutant inflorescences did not include genes that were proposed to be acting downstream of *TAW1* such as *SVP* family MADS-box genes or *OsMADS7, OsMADS8, OsMADS16, OsMADS3* and *OsMADS58* (Yoshida et al., 2013). This observation suggests that, as being part of two distinct phylogenetic groups, *OsG1L2* (and probably also *OsG1L1*) may function in an inflorescence developmental pathway that acts in parallel with *TAW1*. However, further analysis will be needed to sustain this hypothesis since the DEG list published by Yoshida et al. (2013) was obtained from microarray studies using the *taw1-D* gain-of-function mutant whereas we used the *osg1l2* knock-out mutant which showed an opposite phenotype.

The transcriptomic analysis of the *osg1l2* reproductive meristems revealed that some of the differentially expressed genes are known to be involved in inflorescence architecture determination. For instance, in previous studies, *OsRCN1* and *OsRCN4* overexpression resulted in plants that produced panicles with an increased number of branches (Nakagawa et al., 2002; Liu et al., 2013) and knocking-down all RCN genes resulted in shorter panicles with fewer secondary branches (Liu et al., 2013). This phenotype is similar to the one observed in the *osg1l2* mutant, where *OsRCN1* and *OsRCN4* are both downregulated. Furthermore, Nakagawa et al. (2002) indicate a role of *RCN* genes in the suppression of spikelets meristem identity.

Another example is *OsGATA7* which was downregulated in the *osg1l2* mutant. Knock-down and knock-out mutants showed alterations in the architecture of the inflorescence. Knock-down of *OsGATA7* results in panicles with a reduced number of primary branches, whereas the knock-out mutant develops panicles bearing fewer primary and secondary branches (Zhang et al., 2018). Based on the results obtained from our analyses it could be suggested that deregulation of *OsRCN1, OsRCN4* and *OsGATA7* expression in the *osg1l2* mutant might contribute to the observed inflorescence phenotypes. Since in the predicted GRN, *OsGATA7* resulted to be located in a different subcluster than *OsG1L2* (Figure 4), it is tempting to speculate that *OsGATA7* regulates inflorescence architecture in a parallel pathway.

In the GRN proposed, *PAP2/OsMADS34* falls in the same subcluster as *OsG1L2*. OsMADS34 belongs to the *SEPALLATA* (*SEP*) subfamily of the MADS-box gene family and is one of the key regulators of spikelet meristem identity in rice. However, for *pap2/osmads34* mutants contrasting phenotypes have been published. Kobayashi et al. (2010) observed in the *pap2-1* mutant a suppression of the transition of branch meristems into spikelet meristem identity resulting in more primary and secondary branches. While the *osmads34* mutant described by Gao et al. (2010) showed also more primary branches but less secondary branches and spikelets. Both mutants had smaller panicles. Interestingly, in the *osg1l2* mutant, *OsMADS34* expression was reduced, which correlates with the observed reduced panicle length and the reduced number of SBs and spikelets observed in the *osmads34* mutant described by Gao et al. (2010). However, the reduced number of PBs in the *osg1l2* contrasts with the *OsMADS34* downregulation. Overall, *OsMADS34/PAP2* seems predominantly involved in promoting spikelet identity whereas *OsG1L2* plays most likely an opposite role in this transition. Since *OsMADS34* is still expressed in the *osg1l2* mutant it is difficult to conclude which genetic interactions these genes may have. However, it is likely that their pathways are connected, and the balance of their activities may determine the final architecture of the panicle. The relationship between *OsMADS34/PAP2* and *OsG1L2* was furthermore predicted by the GRN that we developed in this study (Figure 4).

Some genes that were upregulated in the *osg1l2* background had an expression level equal to zero in the wild type background. One of them is *OsESP*, which encodes for a long non-coding RNA (Luan et al., 2019). In the semi-dominant *Epi-sp* mutant, 3’ region of the transcribed region was characterised by a loss of DNA methylation resulting in a strong upregulation of the gene causing the development of a denser and shorter panicle. It is possible that the reduction in panicle length observed in *osg1l2* background is linked to the observed upregulation of *ESP*. It would be interesting to investigate whether the loss of *OsG1L2* activity leads to changes in the 3’ methylation of the *ESP* gene. *ESP* was one of the predicted direct targets of OsG1L2 within the GRN. The GRN also indicated that *ESP* (together with other 8 genes, still unknown and only expressed in *osg1l2* mutant developing inflorescences) is a target of both *OsG1L2* and *OsFC1*. It is known that *OsFC1* controls panicle architecture since the null mutant developed smaller panicles (Cui et al.,2020). It is tempting to hypothesize that OsFC1 during the reproductive phase together with OsG1L2 represses *ESP*. Among the genes which we selected for real-time PCR validation, *OsHOX14* showed a strong downregulation, especially in the *osg1l2* mutant background. OsHOX14 is a member of the homeodomain-leucine zipper (HD-Zip) transcription factor family, and is known to play important roles in different aspects of plant development and morphogenesis and also in responses to biotic and abiotic stresses (Sessa et al., 2018). In particular, the results obtained in different species suggest a role of some HD-Zip I family proteins as integrators of internal and external signals in the regulation of abiotic and biotic stresses as well as in specific growth and developmental pathways (Perotti et al., 2017). The specific expression profile obtained through in-situ hybridization, confirmed the expression of *OsHOX14* in reproductive meristems as reported by Harrop et al. (2016) and Shao et al. (2018). It is furthermore worth mentioning that the online RiceXPro tool (https://ricexpro.dna.affrc.go.jp/) showed the expression of *OsHOX14* mainly in developing panicles and pistils, suggesting a specific role of this transcription factor during reproductive development. The CRISPR *oshox14* mutants that we generated, confirmed a role for this gene in inflorescence development, since panicle development was impaired in these mutants, carrying fewer PBs and SPs when compared to wild-type plants. This phenotype is similar to the *g1l2* mutant in which *OsHOX14* expression was strongly reduced. The *OsHOX14* overexpression lines generated by Shao et al. (2018) displayed dramatic phenotypes, such as severe delay in growth at the seedling stage and difficulties with panicle exertion through stem and leaves. A low-overexpression line was analysed for panicle development which showed both a reduction in panicle length and PB number. This is coherent to our observations in the *oshox14* knock-out CRISPR mutant. Since ectopic expression of *OsHOX14* during the vegetative phase causes growth defects it might well be that this has pleiotropic effects on reproductive development. However, we cannot rule-out the possibility that the knock-out mutant caused a similar phenotype as the overexpression line because of the existence of regulatory loops and dependence on threshold levels that could influence regulatory pathways (Prelich, 2012).

Comparing the *osg1l1* and *osg1l2* CRISPR mutants revealed that the *osg1l2* mutant rice plants present a more severe reduction of SBs. Furthermore, the *osg1l2* panicles also present fewer spikelets than the *osg1l1* single mutant. For these reasons, we hypothesize that *OsG1L2* plays a more important role than *OsG1L1* in inflorescence architecture development. Based on the RNA-seq data, it is unlikely that *OsG1L1* is regulated by OsG1L2, since *OsG1L1* was not deregulated in the *osg1l2* background. However, it is very well possible that *OsG1L1* and *OsG1L2* act in the same pathway. This hypothesis is further supported by our qRT-PCR studies that showed that the expression of some genes was deregulated in both mutants.

Overall, our analysis suggests that *OsG1L1, OsG1L2* and *TAW1/OsG1L5* seem to act in none or partially overlapping pathways. Furthermore, we propose for *OsG1L2* and *OsG1L1* a role in IM specification, branch formation, spikelets number determination and in seed development. Moreover, *OsG1L2* seems to act in pathways that include *OsMADS34, OsHOX14* and *OsFC1*. Finally, the functional analysis of *OsHOX14* indicates that the proposed GRN promises to be of value for the identification of new players in the first stages of inflorescence development.

## MATERIALS AND METHODS

### 1. Plant material and growth condition

For the experiments we used *Oryza sativa, ssp. japonica, cv Nipponbare*. The plants were grown for 8-10 weeks in LD conditions (70% humidity, 16h light at 28°C/8h dark at 26°C) and then moved in SD conditions (70% humidity, 12h light at 28°C/12h dark at 26°C) to induce flowering. In vitro, plants were germinated on MS-F medium (2,2 g/L MS + vitamins, 15 g/L Sucrose, 1L ddH2O, pH adjusted to 5.6 adding KOH, 2.5 g/L gelrite) and after 15 days were transplanted in soil. Plants used for phenotypic analysis were grown in IRD transgenic greenhouse (Montpellier, France) in spring 2019 (seeds have been sown in February 2019 and panicles collected in June) under natural day conditions at 28°C-30°C, and humidity at 60%. Plants used for the RNAseq experiment were grown in the C chamber in NC State University Phytotron.

### 2. RNA isolation and cDNA synthesis

Total RNA from different tissues (roots, young and mature leaves, milk and mature seeds) and from meristematic tissue enriched in Inflorescence Meristems (IM), Primary Branch Meristems (PBMs), Elongated Primary branch meristems and Axillary Meristems (ePBM/AM) and Spikelet Meristems (SM) was extracted with the NucleoSpin® RNA Plant kit (http://www.mn-net.com) and DNA contamination was removed using the TURBO DNA-free™ Kit according to the manufacturer’s instructions (https://www.thermofisher.com). The different tissues were sampled in liquid Nitrogen using an optical microscope. The RNA was reverse transcribed using the ImProm-II™ Reverse Transcription System (https://ita.promega.com) and the cDNA was used as a template in RT-PCR reactions.

### 3.1 Transcriptome analysis

80 plants (40 WT plants and 40 *osg1l2* mutant plants) were sown in growth chamber under LD condition (70% humidity, 16h light at 28°C/8h dark at 26°C), at NC State University Phytotron and, after 12 day of induction in SD conditions, were sampled. Fifty mg of tissue, corresponding to 8-10 meristems at early developmental stages enriched in PBMs and ePBM/AMs, was manually dissected using an optical microscope. RNA was extracted from the samples using RNeasy Plant Mini Kit from Qiagene. cDNA libraries were prepared using NEBNext Ultra DNA Library Prep Kit for Illumina (E7370) according to the manufacturer’s instructions. Novaseq6000 Illumina machine was used and sequencing was single-end stranded.

### 3.2 Data analysis

Raw RNA-seq data in fastq format of wild type and *osg1l2* mutant was processed and subsequently used for gene regulatory network inference with the TuxNet interface (Spurney et al., 2019) (https://github.com/rspurney/TuxNet). For the processing of the raw RNA-seq, the gff and fasta file of the reference genome (IRGSP-1.0) and gene name file was downloaded from The Rice Annotation Project Database (RAP-DB; https://rapdb.dna.affrc.go.jp/download/irgsp1.html) (Supplemental Table S5). The gene IDs from rice transcription factors were downloaded from the plant transcription database v4.0 (http://planttfdb.gao-lab.org/) and converted from MSU to RAP, with the manual addition of the *ALOG* gene family. Next, TuxNet uses ea-utils fastq-mcf (Aronesty, 2013) for pre-processing, hisat2 (Kim et al., 2015) for genome alignment and Cufflinks (Trapnell et al., 2012) for differential expression analysis. The following xlsx files are generated by TuxNet: a file containing the FPKM values for each replicate (Supplemental Table S3), a file with the differentially expressed genes (DEGs) identified with a q-value threshold of 0.05 and a log_2_(fold change) of 1 (Supplemental Table S4), and a file containing an average expression, a log_2_(fold change), and a q-value (Supplemental Table S6).

The PCA (principal component analysis) was performed in R with the RPKM file from TuxNet using the *prcomp* function from the *stats* package and the *pca3d* package (January Weiner (2020). pca3d: Three Dimensional PCA Plots. R package version 0.10.2. https://CRAN.R-project.org/package=pca3d) (Supplemental Figure S5).

To infer the gene regulatory network (GRN) in *osg1l2* and predict the causal relationships between and target genes underlying the inflorescence phenotype in *osg1l2*, the differentially expressed TFs and genes identified in *osg1l2* with a q-value threshold < 0.05 and a log_2_(fold change) > 1 or < −1 (Supplemental Table S4) were selected. We manually added *OsG1L2* to the DEGs list, since the log_2_FC of *OsG1L2* was less than 1 (−0.885798). Within the TuxNet interface, RTP-STAR (Regression Tree Pipeline for Spatial, Temporal and Replicate data) leverages the replicate data of the wild type and *osg1l2* and consists of three parts: spatial clustering using the k-means method, network inference using GENIE3 (regression tree with random forest approach) and edge sign (activation or repression) identification using the first-order Markov method. As options, we used 100 iterations when inferring the GRN and an edge proportion equal to 0.33. The table containing the final predicted network (Supplemental Table S7) has been imported into Cytoscape® 3.8.0 (Shannon et al., 2003), a network visualization software, to obtain high-quality graphics representation of the predicted GRN. Different node shape, color, and size were used to represent TFs, the down- or upregulation in *osg1l2*, and the number of interactions, respectively. The nodes within the network were clustered into different modules with the Cytoscape plugin clusterMaker2 according to the available community clustering algorithm, an implementation of the Girvan-Newman fast greedy algorithm that uses connectivity to cluster nodes (Morris et al., 2011).

### 4. qRT-PCR Analysis

Fresh meristematic tissue enriched in Inflorescence Meristems (IM), Primary Branch Meristems (PBMs), Elongated Primary branch meristems and Axillary Meristems (ePBM/AM) and Spikelet Meristems (SM) was collected in liquid nitrogen using an optical microscope. RNA was extracted with the NucleoSpin® RNA Set for NucleoZOL - MACHEREY-NAGEL KIT for high purity products. The qRT-PCR analysis was carried out in a final volume of 12 μL in a Biorad C1000™ thermal cycler, using 3 μL of a 1:10 dilution cDNA, 0,2 μM (stock 10mM) Forward and Reverse Primer, 6 μL of Sybr Green Super Mix 2X (Bio-Rad), 2,6 μl MQ H2O.

The expression levels of *OsG1L1(LOC_Os02g07030), OsG1L2 (LOC_Os06g46030)* and *TAW1 (LOC_Os10g33780)* were evaluated using primer pairs RT2541/RT2542, RT1387/ RT1389 and RT2543/ RT2544 respectively. The RT-PCR was performed with the following conditions: 95°C 90’’ 40 cycles (95°C 15’’, 60°C 10’’, 60°C 30’’) and 60°C 10’’.

The expression levels of *OsESP(Os01g0356951), OsMADS34/PAP2 (LOC_Os03g54170), OsHOX14 (LOC_Os07g39320), OsCEP6 (LOC_Os08g37070*.*1), OsTB1/FC1 (LOC_Os03g49880)* and *Os03g0569000* in WT and *osg1l2* background were evaluated using primer pairs OSP2055/OSP2056, OSP0855/OSP0856, OSP1400/1401, OSP2043/OSP2044, OSP2045/OSP2046, OSP2059/OSP2060 and using the following condition: 95°C 90’’ 40 cycles (95°C 15’’, 58°C 10’’, 60°C 30’’) and 60°C 10’’.

Three biological replicates for each experiment were performed.

Rice *Elongation Factor 1* (*EF1*) (*LOC_Os03g08010*) was used as an internal reference during the experiments. Primer sequences are listed in Supplemental Table S8.

### 5. Tissue fixation and *In situ* Hybridization

Rice reproductive meristems from the main stem at different stages of early panicle development were collected and fixed in FAA [ethanol (Fluka) 50 %; acetic acid (Sigma-Aldrich) 5 %; formaldehyde (Sigma-Aldrich) 3·7 % (v/v)], infiltrated under mild vacuum conditions for 15 min in ice. After 1h 45’ the samples were washed 3 times for 10’ in EtOH 70% and conserved at 4°C; they were dehydrated in a series of increasing graded ethanol series, transferred to bioclear (Bioptica) and then embedded in Paraplast X-TRA® (Sigma-Aldrich). To generate the sense and antisense probes, gene fragments were amplified from cDNA using gene-specific primers (Supplemental Table S8), cloned into pGEM®-T Easy Vector and confirmed by sequencing. Digoxigenin-labeled antisense and sense RNA probes were transcribed and labelled from pGEM^®^-T Easy with T7/SP6 RNA polymerase (Promega) according to the manufacturer’s instructions and using the DIG RNA labelling mix (Roche). Paraplast-embedded tissues were sliced on an RM2155 microtome (Leica) at 8 μm of thickness and hybridized as described by Caselli et al. (2019) with minor modifications. Immunodetection was carried out with anti-digoxigenin-AP Fab fragment (Roche) and BCIP-NBT colour development substrate (Promega) as specified by the manufacturer. Sample’s images were acquired with a Zeiss Axiophot D1 (Zeiss, Oberkochen, Germany) microscope with an Axiocam MRc 5 (Zeiss) at different magnifications.

### 6. CRISPR-Cas9 construct generation

For the generation of *osg1l1, osg1l2* and *oshox14* (LOC_Os07g39320) single knock-out mutants, 20-bp specific protospacers (Supplemental Table S8) for each gene were selected using the CRISPR-P database (http://cbi.hzau.edu.cn/crispr/) and cloned into the *BsaI* site of pOs-sgRNA entry vectors under U3 promoter and then combined into the destination vector containing the Cas9 under maize Ubiquitin Promoter using the Gateway® LR Clonase II Enzyme mix following the procedure reported by Miao et al. (2013) and already followed by Lacchini et al., 2020.

### 7. Bacterial and plant transformation

For bacterial transformation, we used *E. coli* electrocompetent cell (DH10b strains) and *Agrobacterium tumefaciens* electrocompetent cell (EH105 strain).

All final constructs were used to transform embryogenic calli obtained from *Oryza sativa* L. ssp. *japonica cv. Nipponbare* seeds according to the methods described by Hiei et al. (1994) and Toki (1997).

### 8. Mutant screening in transgenic plants

Genomic DNA was extracted from T0-hygromycin-resistant rice plants and genotyped by PCR using primers specific for Cas9 construct, Atp5706/Atp5718 (Supplemental Table S8). Subsequently, from the positive plants, DNA fragments across the target sites were amplified through PCR using gene-specific primer pairs (Supplemental Table S8). The PCR amplicons were purified and sequenced. The obtained chromatograms were analysed and compared with WT sequences with FinchTV searching for mutations.

### 9. Phenotypical analysis of panicles and seeds

To perform phenotypical analysis 15, 19 and 20 panicles from the main tiller were collected respectively from WT, *osg1l1* and *osg1l2* plants. Each panicle was attached on A4 white paper and all panicle branches were spread and blocked with transparent sticks. Each paper with panicle and scale bar was put on an Image capturing system consisting of Portable Camera Stand and two RB 218N HF Lighting Units. The pictures were processed into P-TRAP software. The analysis was done as described in AL-Tam et al., (2013). The results were statistically analysed by One Way ANOVA followed by Tukey test and represented with GraphPad Prism 8.

To perform the phenotypical analysis on wild type and *oshox14* plants, all the panicles produced by 5 wild-type and 5 *oshox14* plants were collected. For each panicle, the number of Primary Branches (PBs), Secondary Branches (SBs) and Spikelets/seeds (SP) was manually calculated. The results were statistically analysed with a Student’s T-test and graphically represented with GraphPad Prism 8.

At least one hundred seeds were analysed for each genotype (*osg1l1, osg1l2* and WT). Images of the seeds were acquired using a Leica MZ6 stereomicroscope in conjunction with a Leica DFC280 camera at different magnifications; each image was then processed with Smart Grain (Tanabata et al. 2012). The obtained results were statistically analysed with a One Way ANOVA followed by Tukey test and represented with GraphPad Prism 8.

## Accession Numbers

Sequence from this article can be found in the GeneBank / EMBL databases under the following accession numbers: *OsG1L1 LOC_Os02g07030, OsG1L2 LOC_Os06g46030, TAW1 LOC_Os10g33780, OsESP Os01g0356951, OsMADS34/PAP2 LOC_Os03g54170, OsHOX14 LOC_Os07g39320, OsCEP6 LOC_Os08g37070.1, OsTB1/FC1 LOC_Os03g49880, Os03g0569000, OsEF1 LOC_Os03g08010*

## Supplemental Data

The following supplemental materials are available:

**Supplemental Figure S1** Gene structure and wild type and mutant proteins alignment of *OsG1L1*.

**Supplemental Figure S2** Gene structure and wild type and mutant proteins alignment of *OsG1L2*.

**Supplemental Figure S3** Chromatogram showing the type of mutation and phenotypical analysis of *osg1l2* mutant (C insertion).

**Supplemental Figure S4** Gene structure and wild type and mutant proteins alignment of *OsHOX14*.

**Supplemental Figure S5** PCA output.

**Supplemental Table S1** Phenotypical traits analysed in wild type, *osg1l1* and *osg1l2* plants.

**Supplemental Table S2** Area, length and width of wild type, *osg1l1* and *osg1l2* seeds.

**Supplemental Table S3** FPKM values for each replica.

**Supplemental Table S4** Differentially expressed genes between wild type and *osg1l2* mutants.

**Supplemental Table S5** List of gene names.

**Supplemental Table S6** Average gene expression, log_2_(fold change) and q-value in wild type and *osg1l2* mutant.

**Supplemental Table S7** Final predicted GRN.

**Supplemental Table S8** Primers used in this article.

## ACKNOWLEDGMENTS

We like to thank Andrea Guazzotti and Marco Maffei for their help with the analysis of the *oshox14* mutant. Furthermore, we thank Mario Beretta and Valerio Parravicini for taking care of the rice plants.

## Notes

1 This work was supported by MUR PRIN2017_2017N5LBZK The PhD fellowship of E.F. and V.M.B, were supported by the Doctorate School in Molecular and Cellular Biology, Università degli Studi di Milano. E.F. was supported by H2020-MSCA-RISE-2015 ExpoSEED Proposal Number: 691109.

